# Impact of air pollution on Neurite Orientation Dispersion and Density metrics (NODDI) in 10–13-year-old children with and without ADHD diagnosis

**DOI:** 10.1101/2025.11.03.686321

**Authors:** Paulina Lewandowska, Claude J. Bajada, Mikołaj Compa, Yarema Mysak, Aleksandra Domagalik, Bartosz Kossowski, Clemens Baumbach, Katarzyna Kaczmarek-Majer, Anna Degórska, Krzysztof Skotak, Katarzyna Sitnik-Warchulska, Małgorzata Lipowska, Bernadetta Izydorczyk, James Grellier, Iana Markevych, Marcin Szwed

## Abstract

Air pollution is a significant risk factor for adverse neurodevelopmental outcomes in children. While studies have linked pollutants to changes in brain structure, specific effects on white matter microstructure remain inconclusive. Neurite Orientation Dispersion and Density Imaging (NODDI) provide two nuanced, separate white matter measures that serve as proxies for neurite density and cell-body organization. We used NODDI to examine potential associations between prenatal, early life and current exposure to nitrogen dioxide (NO_2_) and particulate matter with diameter < 10 micrometers (PM_10_) and white matter microstructure in school-aged children. We also explored whether ADHD diagnosis moderated these associations. We observed several negative associations between both NO_2_ and PM_10_ exposure and neurite density across various white-matter fibers and exposure windows, but none of the associations were statistically significant after correcting for multiple comparisons.

In this analysis, we did not find any statistically significant associations between long-term exposure to PM_10_ and NO_2_ and white matter microstructural integrity as measured by NODDI. Our results highlight the challenge of detecting modest environmental impacts on the brain and underscore the need for larger, more targeted studies to confirm these preliminary trends.

## 1. INTRODUCTION

Air pollution is one of the leading environmental contributors to the disease burden worldwide (Global Burden of Disease Collaborative Network, 2024). Exposure to airborne pollutants, such as fine particulate matter (PM) and nitrogen dioxide (NO_2_) has been consistently linked to respiratory (Duan et al., 2020; Paciência et al., 2022) and cardiovascular (de Bont et al., 2022; Konduracka & Rostoff, 2022) diseases. A growing body of research has highlighted the effects of air pollution on the nervous system, particularly exposure during critical periods of childhood development (Cory-Slechta et al., 2023; Herting et al., 2024; Morrel et al., 2025; Parenteau et al., 2024). Children are especially vulnerable to air pollution due to greater effective exposure from physiological and behavioral factors, such as higher inhalation volume per body weight (Bateson & Schwartz, 2008). This heightened biological sensitivity means that as pollutants cross the blood-brain barrier, they can disrupt the maturing brain via mechanisms like neuroinflammation and microglial activation, leading to potential long-term consequences (Cory-Slechta et al., 2023; Morris et al., 2021). Consistent with these mechanisms, childhood exposure to air pollution has been linked to cognitive deficits, particularly in attention and executive functioning (Compa et al., 2023; Forns et al., 2016; Sunyer et al., 2015). This link is further solidified by growing evidence connecting early life exposure to traffic-related air pollution with an increased risk of Attention Deficit Hyperactivity Disorder (ADHD), a neurodevelopmental condition characterized by these specific impairments (Liu et al., 2023; Thygesen et al., 2020).

Although growing evidence suggests that air pollution can affect cognitive development and behavior, our understanding of the underlying neural pathways remains limited (Cory-Slechta et al., 2023; Herting et al., 2024; Parenteau et al., 2024; Polemiti et al., 2024; Szwed et al., 2025). Studies employing structural Magnetic Resonance Imaging (MRI) indicate that exposure to pollutants such as particulate matter with an aerodynamic diameter of less than 2.5 micrometers (PM_2.5_) and with a diameter of less than 10 micrometers (PM_10_), NO_2_, and polycyclic aromatic hydrocarbons (PAHs) (Mortamais et al., 2017; Yang et al., 2025) are associated with changes in cortical volume, thickness, and surface area (Guxens et al., 2018; Herting et al., 2019; Kusters et al., 2025; Pujol, Martínez-Vilavella, et al., 2016). However, findings are inconsistent: the reported effects are often small, and the specific cortical or subcortical regions implicated differ substantially across studies (Morrel et al., 2025). In addition to analyses of structural MRI, a small but growing body of research has used resting-state functional MRI (rs-fMRI) to explore the impact of air pollution on brain network organization (Cotter et al., 2023; López-Vicente et al., 2025; Pujol, Fenoll, et al., 2016; Zundel et al., 2024). Most recent longitudinal studies are beginning to shed light on how these exposures affect the maturation of brain networks over time (Cotter et al., 2023; López-Vicente et al., 2025). The majority of studies reporting associations between air pollution exposure and altered white matter have relied on metrics from Diffusion Tensor Imaging (DTI) (Binter et al., 2022; Kusters et al., 2024; Lubczyńska et al., 2020; Pujol, Martínez-Vilavella, et al., 2016). While the majority of such studies typically report statistically significant associations between air pollution exposures and altered white matter (Binter et al., 2022; Burnor et al., 2021; Kusters et al., 2024; Lewandowska et al., 2025; Lubczyńska et al., 2020; Pujol, Fenoll, et al., 2016; Pujol, Martínez-Vilavella, et al., 2016), their findings are limited by a lack of biological specificity. DTI metrics like fractional anisotropy (FA) are informative, they provide only limited insight into the precise microstructural properties of the brain tissue; for example, FA conflates multiple features such as neurite density, myelination, and fiber organization, leading to potentially ambiguous interpretation (Jones et al., 2013). More advanced diffusion models, such as neurite orientation dispersion and density imaging (NODDI), restriction spectrum imaging (RSI), and fixel-based analysis (FBA), offer a more nuanced characterization of white matter microstructure compared to a conventional tensor model (Burnor et al., 2021; Cotter et al., 2024). While studies on the Adolescent Brain Child Development (ABCD) cohort have successfully used RSI to identify microstructural alterations linked to air pollution (Burnor et al., 2021; Cotter et al., 2024), our own previous work on the Neurosmog cohort found no significant associations using FBA (Lewandowska et al., 2025). This discrepancy suggests either that the effects are different in the two cohorts, or that the detection of any underlying biological effect is sensitive to the specific microstructural properties captured by different diffusion models. To address this issue, the present study employs NODDI, an advanced method of diffusion modelling, not yet applied in this research context. It has already been used in developmental research to track brain maturation in children (Lynch et al., 2020; Mah et al., 2017; Vaher et al., 2022) and proved to be a sensitive metric of microstructural changes across early life. However, its application in the field of environmental neuroimaging remains scarce. Conducting such research is particularly important as evidence indicates that exposure to air pollution can increase the risk for the incidence of ADHD and its associated symptoms (Bernardina Dalla et al., 2022; Liu et al., 2023).

In the present study we therefore applied NODDI to examine potential associations between air pollution exposure and white matter microstructure in children with ADHD diagnosis and their typically developing peers. We hypothesized that higher exposure to air pollution would be associated with altered neurite density and orientation dispersion in atlas-based white matter regions. Guided by our prior work on the same population (Lewandowska et al., 2025), we focused on NO_2_ and PM_10_ exposure during the first four years of life and conducted an additional analysis of the prenatal period (Binter et al., 2022; Bové et al., 2019; Crooijmans et al., 2024). As an exploratory step, we also examined associations between NODDI metrics and current long-term exposure. Prior studies have highlighted the potential links between air pollution exposure and attentional deficits (Compa et al., 2023; Liu et al., 2023; Thygesen et al., 2020) suggesting that there is a possibility that both factors may independently or jointly influence brain structure. Additionally, structural brain differences between children with and without ADHD in both white and grey matter have been reported (Bernanke et al., 2022; McAlonan et al., 2007; Nakao et al., 2011). Also, given that ADHD is linked to baseline structural alterations in white matter (Connaughton et al., 2022; Fuelscher et al., 2021) and may represent a vulnerability factor, we also hypothesized that ADHD diagnosis moderates these associations.

## 2. MATERIALS & METHODS

Our analysis was preregistered on the Open Science Framework (OSF) registry as a project entitled “Impact of prenatal and early life exposure of air pollution on NODDI metrics in Polish children: diffusion MRI study.” (Lewandowska, 2025). We used data from the NeuroSmog study (Markevych et al., 2021), which was conducted according to the guidelines of the Declaration of Helsinki and approved by the Ethics Committee of the Institute of Psychology, Jagiellonian University, Kraków, Poland (#KE_24042019A). Written informed consent was obtained from the legal guardians of all children enrolled in the study, and all children also gave their own written informed assent. The Clinical Trials Identifier is NCT04574414.

### 2.1 Study population and sample size

The final sample consisted of 388 children from the NeuroSmog cohort (Markevych et al., 2021). The children were aged 10–13 years (mean age = 11.3, SD = 0.78; 161 female) and either had a clinical diagnosis of ADHD or were typically-developing (TD) children. In the NeuroSmog study, the children in the TD group were defined as participants without an ADHD diagnosis, intellectual disability, or any known neurological, comorbid psychiatric, or other serious medical conditions (Markevych et al., 2021). The TD children were recruited via random stratified sampling from schools in 18 towns. Study towns were selected based on varying population sizes and air pollution levels. In parallel, children with an ADHD diagnosis were recruited from the same 18 towns using convenience sampling, with referrals from local psychological counseling centers, school psychologists, and medical professionals, or through direct parental application. From the 741 children enrolled in the NeuroSmog study (demographic details in Supplementary Materials, *Supplemental Methods 1*, Table 1S), 714 children completed the MRI scanning session. Children with a complete diffusion-weighted imaging sequence consisted of 578 and these images underwent rigorous quality control processes. After completion of all quality control, preprocessing and registration steps (see section 2.4 *Diffusion data preprocessing and quality control*), the final sample of 388 participants was established (Table 1). For a comprehensive description of the inclusion/exclusion criteria and recruitment details, please refer to the NeuroSmog protocol paper and previous published analyses (Compa et al., 2023; Markevych et al., 2021).

**Table 1.** Descriptive statistics of the study sample.

| Variable |  | ADHD children | TD children | All |
| --- | --- | --- | --- | --- |
|  |  | (n = 109) | (n = 279) | (N = 388) |
|  |  | Mean (SD) | Mean (SD) | Mean (SD) |
| NO<sub>2</sub> (µm) | NO<sub>2</sub> prenatal | 23.5 (9.7) | 23.9 (8.7) | 23.8 (9.0) |
|  | NO<sub>2</sub> in years 0-4 | 24.1 (7.8) | 24.6 (7.0) | 24.4 (7.2) |
|  | NO<sub>2</sub> in current exposure | 19.5 (8.3) | 20.2 (7.6) | 19.98 (7.8) |
| PM<sub>10</sub> (µm) | PM<sub>10</sub> prenatal | 47.2 (16.2) | 44.5 (14.1) | 45.24 (14.8) |
|  | PM<sub>10</sub> in years 0-4 | 44.8 (7.1) | 45.2 (6.4) | 45.07 (6.6) |
|  | PM<sub>10</sub> current exposure | 37.3 (6.6) | 37.2 (5.5) | 37.23 (5.9) |
| Age (years) |  | 11.30 (0.9) | 11.30 (0.2) | 11.3 (0.8) |
|  |  | N (%) | N (%) | N (%) |
| Sex | Female | 29 (26.6) | 133 (47.7) | 162 (41.8) |
|  | Male | 80 (73.4) | 146 (52.3) | 226 (58.3) |

*Note.* ADHD = Attention Deficit Hyperactivity Disorder; TD = Typically Developing; NO<sub>2</sub> = nitrogen dioxide; PM<sub>10</sub> = Particulate matter with diameter less than 10 micrometers; SD = standard deviation.

### 2.2 Air pollution exposure assessment

To estimate individual-level exposure to air pollution, we first collected lifelong residential address histories for each child, provided by their legal guardians (Markevych et al., 2021). For this study, we successfully developed reliable exposure estimates for PM_10_ and NO_2_. Monthly average concentration maps for these pollutants were generated at 125m resolution using hybrid Land Use Regression (LUR) models (de Hoogh et al., 2013; Szwed et al., 2025), as detailed in the Supplementary Materials (Supplemental Methods 2). The prenatal exposure period was estimated by averaging the monthly exposure across the second and third trimester of pregnancy, periods most crucial for fetal brain development. The start of each pregnancy was calculated by subtracting the gestational age from the child’s birth date. For cases where gestational age was not available, we assumed a duration of 40 weeks. Early life exposure corresponds to the average of annual exposure estimated within the first four years of children’s life (later referred to as ‘years 0-4’). Current exposure (years 2020-2022) was assessed by assigning modeled annual average concentrations to each child’s residential address. The models used were based on 2018 monitoring data, as these were the most up-to-date available.

### 2.3 MRI data acquisition

MRI data were acquired at the Małopolska Centre of Biotechnology, Jagiellonian University in Kraków, Poland, using Siemens MAGNETOM Skyra 3T MRI scanner (Erlangen, Germany) with a 64-channel head coil. Children were instructed to remain awake and to stay as still as possible during the scanning process. Each child completed a mock scanning session before the actual scan to familiarize them with the procedure and ensure their comfort. A spin-echo diffusion weighted echo planar imaging (DW_EPI) sequence was applied with TE=101 ms, TR=3800 ms, flip angle = 78 deg, and voxel size = 2 x 2 x 2 mm^3^. We used anterior-posterior (AP) phase coding directions and 118 volumes in total with b-values of 500, 1250, 2500 (n = 18, n = 36, n = 53) along with 11 images applying no diffusion gradient (b-value = 0). To correct for susceptibility-induced distortions, we also acquired 4 images with a b-value = 0 using posterior-anterior (PA) phase coding. For the purpose of registration steps (see *Masks registration to diffusion space* section) we have also used T1-weighted images with a resolution of 1 x 1 x 1 mm^3^, TE=2.88 ms, TR=2500 ms, and flip angle = 8 deg. The acquisition sequences for T1-weighted data were adopted from the ABCD project (Casey et al., 2018). Further information on the neuroimaging data acquisition is incorporated in the NeuroSmog protocol paper (Markevych et al., 2021).

### 2.4 Diffusion data preprocessing and quality control

From the initially recruited participants (N = 741) we obtained complete diffusion-weighted imaging (DWI) data for 578 children (Figure 1). Each scan was visually inspected to identify any structural abnormalities. Twelve participants were excluded due to cerebellar malformations, and an additional 15 were excluded due to image acquisition issues. This resulted in 551 DWI scans included in the preprocessing pipeline. Following, the diffusion images were matched with their corresponding T1-weighted structural scans to ensure proper registration (for detailed description see section Masks registration to diffusion space in the text below).

**Figure 1.**
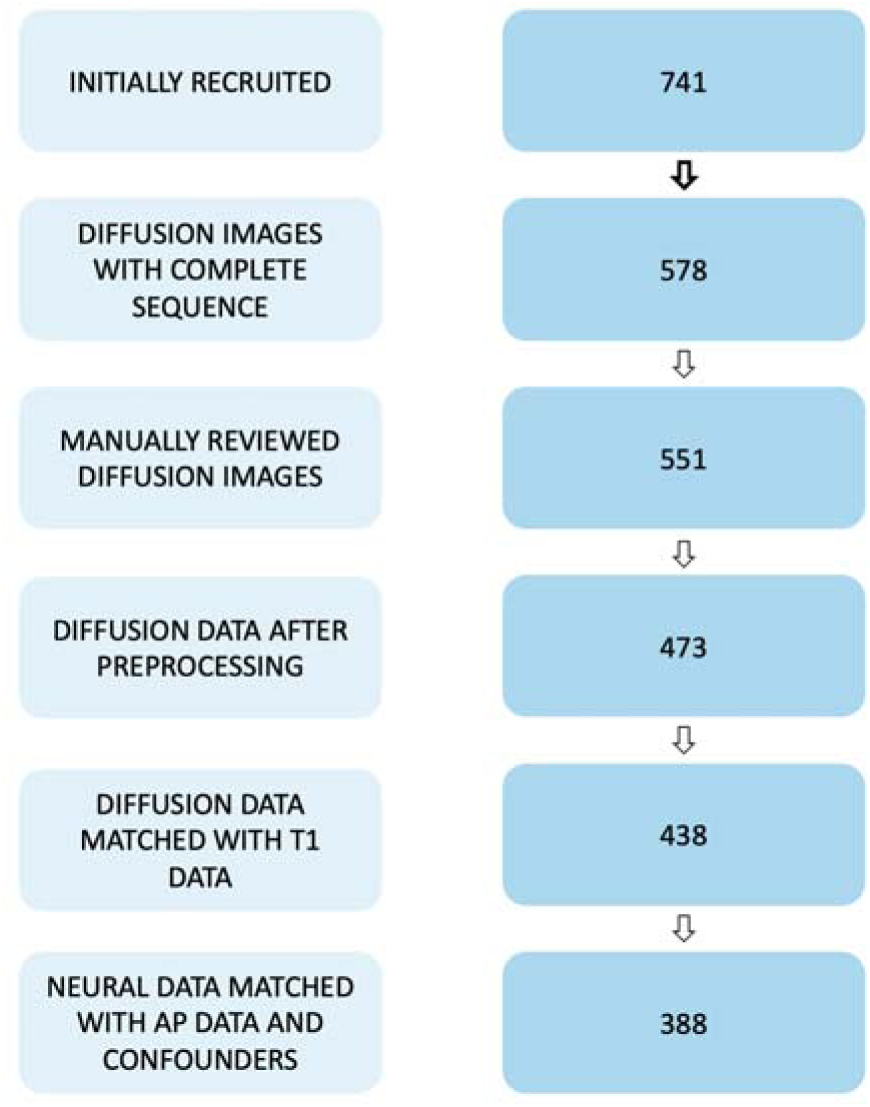
The chart illustrates the number of participants retained at each stage of data processing, starting from initial recruitment to the final sample included in the analyses. Manual review involved excluding images with visible distortions unresolvable during preprocessing or any identified brain malformations. The matching of diffusion with T1 data refers to the co-registration process used to define anatomical regions within the diffusion space. AP: Air Pollution.

Data preprocessing was conducted using the FSL eddy tool (Andersson & Sotiropoulos, 2016) which corrects for subject motion and eddy current-induced distortions. Susceptibility-induced off-resonance fields were estimated from spin-echo EPI images acquired with opposing phase-encoding directions (Andersson et al., 2003), and incorporated into eddy to jointly model susceptibility, motion, and eddy current effects (Andersson & Sotiropoulos, 2016). Slice-level signal loss due to motion during diffusion encoding was identified and replaced using Gaussian Process predictions (Andersson et al., 2016). Intra-volume motion artifacts—wherein misaligned slices compromise volumetric integrity— were corrected using slice-to-volume alignment (Andersson et al., 2017). Subject movement-related variations in susceptibility-induced distortions were modeled using a Taylor series expansion around the first volume of each scan (Andersson et al., 2018).

Quality control (QC) was performed using the eddy QC framework (Bastiani et al., 2019), which provides both subject-level and study-level metrics. Following the study-wise QC protocol implemented in SQUAD, participants with more than one flagged QC index were classified as outliers and excluded from further analysis (n = 78) (see Supplemental Methods 3 for a detailed description of the QC indices).

#### 2.4.1 Selection of white matter ROIs

Due to the limited and inconsistent findings in the existing literature, we adopted a comprehensive, exploratory approach for our region-of-interest (ROI) selection rather than focusing on a small number of pre-selected tracts. To achieve this, we based our analysis on a well-validated, standard-space ICBM-DTI-81 white matter labels atlas (Mori et al., 2005), which parcellates the white matter into distinct tracts and regions. The atlas consists of 81 white matter tracts including brainstem tracts, projection fibers, association fibers, and commissural fibers. The list of the full list of the white matter tracts is available at http://www.bmap.ucla.edu/portfolio/atlases/ICBM_DTI-81_Atlas/ For this study, all available white matter regions from this atlas were selected as our ROIs to provide a thorough and systematic investigation of potential pollution-related effects across the entire white matter architecture.

#### 2.4.2 NODDI estimations

We implemented the NODDI model to gain specific insight into tissue microstructure. NODDI resolves the diffusion signal by modeling three compartments in each voxel: intra-neurite, extra-neurite, and isotropic free water. This approach yields the Neurite Density Index (NDI), which reflects the packing density of axons and dendrites, and the Orientation Dispersion Index (ODI), which quantifies the coherence of neurite orientations, and the Free Water Fraction (FWF), which estimates the proportion of the voxel occupied by non-restricted water, such as cerebrospinal fluid (CSF). Figure 2 provides a schematic illustration of the NODDI decomposition for a representative white matter voxel (Figure 2).

**Figure 2.**
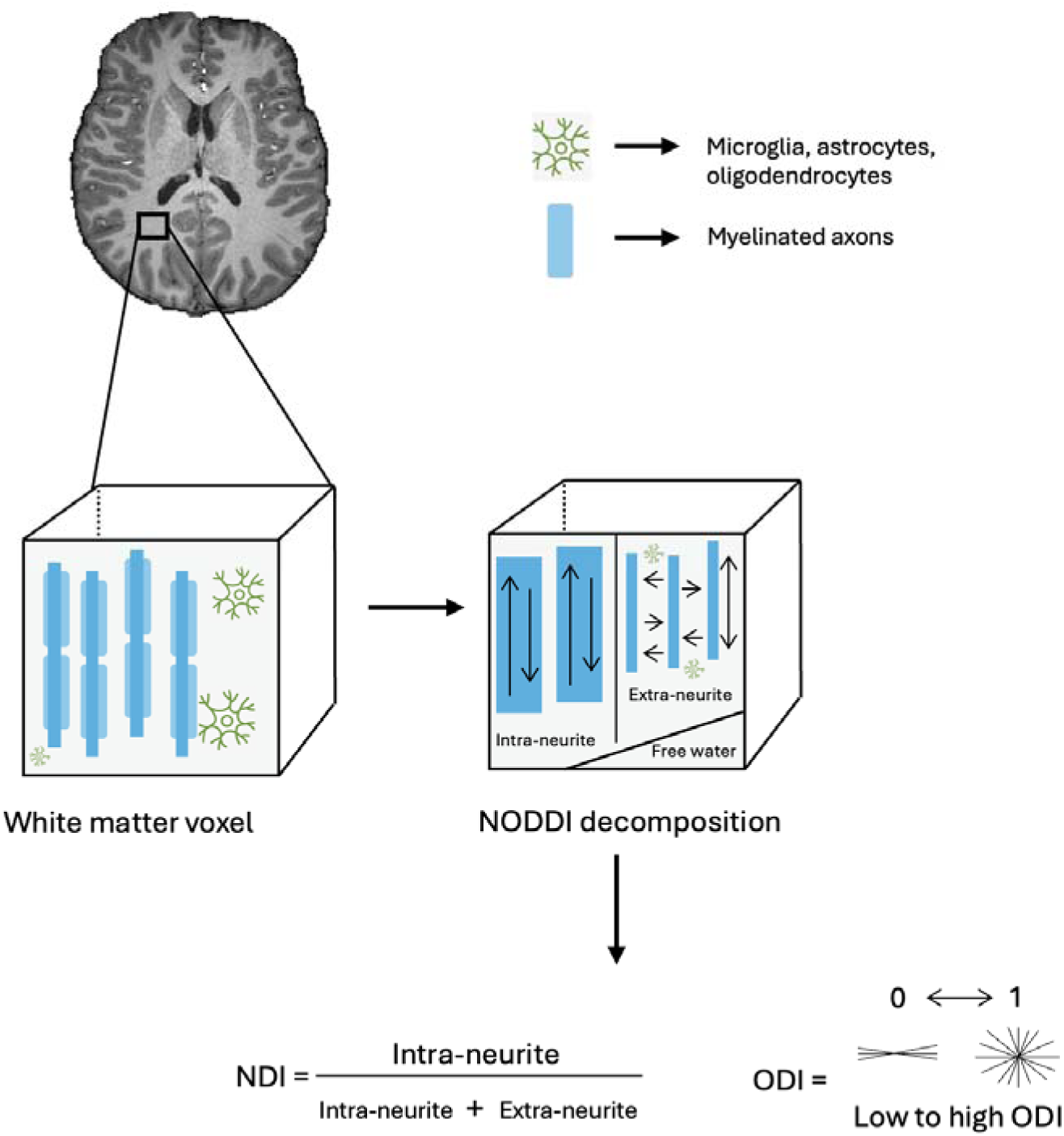
The figure shows how a single white matter voxel is decomposed into three compartments: intra-neurite (water within axons), extra-neurite (water outside axons), and free water. From this, NODDI derives two biophysical metrics: 1) NDI (Neurite Density Index): The volume fraction of the intra-neurite compartment relative to the total tissue volume, defined as the sum of the intra- and extra-neurite spaces (a measure of density); 2) ODI (Orientation Dispersion Index): A measure from 0 to 1 representing the degree of fanning or dispersion of neurite orientation. Low values indicate highly aligned, parallel fibers (e.g., major white matter tracts), while high values indicate complex or crossing fibers (e.g., gray matter).

Estimation of microstructural brain parameters was conducted using the NODDI model (Zhang et al., 2012), implemented via the *CUDIMOT* (Hernandez-Fernandez et al., 2019) toolbox. NODDI parameter estimation was performed using the Watson distribution for modeling specific NODDI metrics. Voxel-wise estimation was carried out using CUDA-accelerated routines with default priors and convergence settings optimized for large-scale application. The model fitting resulted in parameter maps for orientation dispersion and neurite density in native diffusion space. Summary maps were retained for the statistical analyses. All NODDI maps were visually inspected for estimation artifacts and outlier regions prior to ROI extraction within each subject.

#### 2.4.4 Masks registration to diffusion space

For each subject, ROIs were defined using the ICBM-DTI-81 white matter labels atlas (Mori et al., 2005). Then these masks were transformed into each subject’s native diffusion space, as follows. First, the subject’s T1-weighted anatomical image was registered to the standard MNI-152 1mm brain template. This was accomplished using a linear registration with FSL’s *FLIRT* (Andersson et al., 2007; Jenkinson et al., 2012), followed by a non-linear registration with *FNIRT* (Andersson et al., 2007; Jenkinson et al., 2012) to account for individual brain morphology. This process yielded a high-precision warp field from T1 space to MNI space. Next, the inverse of this transformation was applied to each standard-space ROI mask, bringing it into the subject’s native T1 space. The resulting masks were visually inspected against the structural T1 image to confirm accurate alignment. A separate affine registration was then performed using *FLIRT* (Andersson et al., 2007; Jenkinson et al., 2012) to map the skull-stripped T1 image to the subject’s b=0 diffusion image, where the b=0 image was extracted from the first volume of the diffusion-weighted series. Finally, the MNI-to-T1 transformation (derived by inverting the T1-to-MNI warp) and the T1-to-b0 affine transformation were concatenated into a single, combined warp. This transformation was applied to each of the original MNI space ROI masks in a single interpolation step, directly placing the masks into the subject’s native diffusion space. The quality of the final mask alignment was verified by overlaying the binarized ROIs on the diffusion image. NODDI-derived metrics were subsequently calculated within these registered ROIs, ensuring that all analyses were performed in identically defined anatomical regions across subjects. Graphic representation of the mask registration is presented in the Supplementary Materials (Supplemental Methods 4, Figure 2S).

### 2.5 Statistical analysis

All statistical analyses were conducted in R, version 4.4.2. (R Core Team et al., 2024). As a first step, we performed diagnostic checks to assess the distribution of all the variables and to examine the linearity of associations. We verified all statistical assumptions before conducting the analysis. Model residuals were diagnosed using the DHARMa package (Hartig & Hartig, 2020). Based on these diagnostics, we identified non-normally distributed residuals from regression and therefore decided to apply permutation testing to all models using the lmPerm package (Torchiano & Wheeler, 2025), which provides a robust alternative to classical parametric inference. We treated the brain measures as the outcomes (dependent variables) and air pollution as the primary predictor (independent variable). Associations between every dependent variable and the main predictor (air pollution) were assessed by separate models.

We have run all the models based on the formula below:

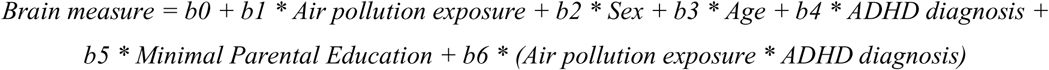

We included an interaction term for air pollution and ADHD status to explore whether the ADHD status moderated the association between air pollution exposure and specific brain outcomes. Exact age in years and sex were included as covariates as they are known to influence brain maturation (Genc et al., 2018; Simmonds et al., 2014). We also included socioeconomic status (represented by minimal parental education with levels low, medium, and high) as a covariate. This represents a deviation from our preregistered analysis plan. We made this decision on the basis of peer-review of a previous manuscript (Lewandowska et al., 2025) which highlighted the importance of controlling for this potential confounder.

## 3. RESULTS

A detailed description and a graphical representation of the NODDI measures are given in Figure 2 in the Methods section (*2.4.2 NODDI estimations*).

### 3.1 Associations between NODDI measures and prenatal and early life air pollution exposures

For NDI, our analysis revealed several main effects for various brain regions in relation to air pollution exposure, which were significant before corrections for multiple comparisons (Table 2). Negative associations were found between NDI and prenatal exposure to NO_2_ in the left cingulum (β = −0.0005, p = 0.028). For early-life exposure (years 0-4), NDI was negatively associated with NO_2_ in the left anterior limb of the internal capsule (β = −0.0004, p = 0.049), the left cingulum (β = −0.0006, p = 0.028), and the right external capsule (β = −0.0005, p = 0.038). Early-life PM_10_ exposure was also negatively associated with NDI in the right anterior corona radiata (β = −0.0006, p = 0.045), left cingulum (β = −0.0007, p = 0.020), right inferior fronto-occipital fasciculus (β = −0.0004, p = 0.016), left posterior limb of the internal capsule (β = −0.0005, p = 0.017), and left superior fronto-occipital fasciculus (β = −0.0009, p = 0.018). None of these associations remained significant after controlling the false discovery rate (FDR) at α = 0.05 using the Benjamini-Hochberg procedure. We found no statistically significant main effects of air pollutants on ODI in these models.

**Table 2.**
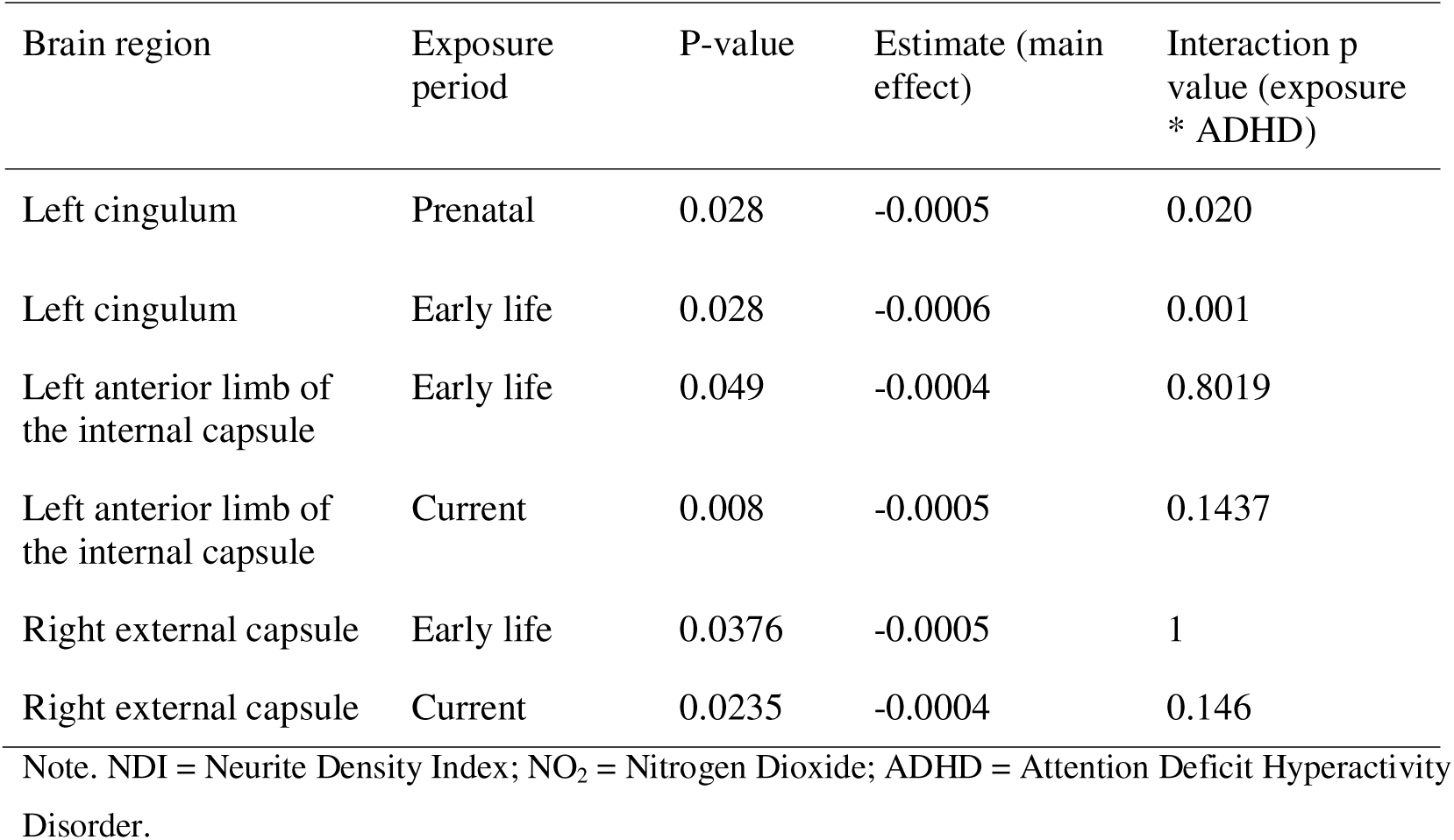
Table shows associations between prenatal and early life exposures to NO_2_ and NODDI measures that were significant at α = 0.05 before multiple-testing correction. The last column shows the p-value for the respective model’s interaction term for NO_2_ and ADHD status.

| Brain region | Exposure period | P-value | Estimate (main effect) | Interaction p value (exposure * ADHD) |
| --- | --- | --- | --- | --- |
| Left cingulum | Prenatal | 0.028 | -0.0005 | 0.020 |
| Left cingulum | Early life | 0.028 | -0.0006 | 0.001 |
| Left anterior limb of the internal capsule | Early life | 0.049 | -0.0004 | 0.8019 |
| Left anterior limb of the internal capsule | Current | 0.008 | -0.0005 | 0.1437 |
| Right external capsule | Early life | 0.0376 | -0.0005 | 1 |
| Right external capsule | Current | 0.0235 | -0.0004 | 0.146 |

Note. NDI = Neurite Density Index; NO<sub>2</sub> = Nitrogen Dioxide; ADHD = Attention Deficit Hyperactivity Disorder.

### 3.2 Additional analysis - associations with current air pollution exposure

Furthermore, negative associations were observed between NDI and current exposure to NO_2_ in the left anterior limb of the internal capsule (β = −0.0005, p = 0.008) and the right external capsule (β = - 0.0004, p = 0.0235) (Table 3). Current exposure to PM_10_ was negatively associated with NDI in the left anterior limb of the internal capsule (β = −0.0006, p = 0.032), the right inferior fronto-occipital fasciculus (β = −0.0005, p = 0.024), and the left superior fronto-occipital fasciculus (β = −0.001, p = 0.015). No significant main effects of air pollutants were found for ODI in these models. None of the associations were statistically significant after controlling the FDR at α = 0.05.

**Table 3.** Table shows associations between prenatal and early life exposures to PM_10_ and NODDI measures that were significant at α = 0.05 before multiple-testing correction. The last column shows the p-value for the respective model’s interaction term for PM_10_ and ADHD status.

| Brain region | Exposure period | P-value (main effect) | Estimate (main effect) | Interaction p value (exposure * ADHD) |
| --- | --- | --- | --- | --- |
| Right anterior corona radiata | Early life | 0.045 | -0.0006 | 0.6348 |
| Left cingulum | Early life | 0.020 | -0.0007 | 0.0176 |
| Right inferior-fronto-occipital fasciculus | Early life | 0.016 | -0.0004 | 0.1669 |
| Right inferior-fronto-occipital fasciculus | Current | 0.024 | -0.0006 | 0.001 |
| Left superior-fronto-occipital fasciculus | Early life | 0.018 | -0.0009 | 0.0849 |
| Left superior-fronto-occipital fasciculus | Current | 0.015 | -0.0011 | 0.096 |
| Left anterior limb of the internal capsule | Current | 0.032 | -0.0006 | 0.0415 |

Note. NDI = Neurite Density Index; PM<sub>10</sub> = Particulate Matter with diameter less than 10 micrometers, ADHD = Attention Deficit Hyperactivity Disorder.

Additionally, several models revealed significant interaction effects between air pollution exposure and ADHD diagnosis, however these were not significant when controlling the FDR at α = 0.05.

## 4. DISCUSSION

We aimed to investigate potential relationships between prenatal, early life, and current exposure to PM_10_ and NO_2_ and white matter microstructure in children aged 10-13. We used NODDI, which allowed for estimating neurite density and orientation dispersion metrics within atlas-based white matter tracts. We found several statistically significant associations between air pollution exposure and the neurite density metric across various white matter regions. However, after correcting for multiple comparisons, neither these associations nor the ADHD moderation analyses remained statistically significant.

To our knowledge, this is the first study to investigate potential associations between air pollution exposure and NODDI metrics referring to intracellular and extracellular spaces. We had hypothesized that air pollution would be associated with localized microstructural alterations, based on previous work using RSI in the ABCD cohort (Burnor et al., 2021). However, our null findings are consistent with our own prior work on the same dataset, which found associations with global DTI metrics but not with the more specific FBA metrics (Lewandowska et al., 2025). This pattern suggests that while global metrics from single-tensor models like DTI may be sensitive enough to detect widespread, subtle effects, the more granular metrics from multi-compartment models like NODDI may require greater statistical power to isolate such changes, especially in a modest-sized sample. Several factors are likely to contribute to this difficulty. First, our final sample (n=388) is substantially smaller than those used in large-scale studies like the ABCD cohort, limiting statistical power. NODDI’s biophysical modeling also required higher data quality standards; specifically, the necessary matching of T1 and diffusion images for accurate registration resulted in a smaller final sample than in our previous DTI/FBA analyses (Lewandowska et al., 2025). Finally, the multiple-comparisons correction required by our broad, atlas-based approach set a high statistical threshold, which may have obscured subtle regional effects.

Although no associations withstood multiple-testing correction, the pattern of the uncorrected findings may offer an exploratory basis for future, more targeted research.

For instance, several uncorrected associations were observed in tracts relevant to attention, which aligns with the proposed neurobiological mechanisms linking air pollution to attentional deficits. Furthermore, the direction of these exploratory associations (e.g., lower NDI with higher exposure) is consistent with RSI findings (Burnor et al., 2021). However, it should be acknowledged that the failure of these associations to survive statistical correction may also reflect a true null finding, indicating no effect. We therefore present these observations not as conclusive evidence, but as exploratory data that could aid in narrowing hypotheses for future studies with more statistical power.

The observed reductions in neurite density within the anterior corona radiata and the left posterior limb of the internal capsule could indicate a potential disruption of large-scale brain networks. For example, the associations in the corona radiata align with previous research linking these areas to the focusing, execution, and shifting of attention, which are all core deficits in ADHD (Stave et al., 2017). The findings in the posterior limb of the internal capsule could further suggest that air pollution may impair critical sensorimotor pathways and broader brain connectivity. Furthermore, the left cingulum, a region with uncorrected associations with prenatal and early-life NO_2_ and early-life PM_10_, is a key component of the default mode network (Burks et al., 2017; Skandalakis et al., 2020) and is proven to be associated with cognitive functioning, such as sustaining attention and working memory (Takahashi et al., 2010). This association between cingulum and air pollution exposure aligns with a previous study exploring air pollution and brain microstructure (Burnor et al., 2021).

Additional uncorrected results pointed to a similar pattern of reduced neurite density associated with both early-life and current air pollution exposure (NO_2_ and PM_10_) in tracts such as the superior fronto-occipital fasciculus (SFOF), the right external capsule (EC), and the left anterior limb of the internal capsule (ALIC). A similar negative association was observed between neurite density and both early-life and current PM_10_ exposure in the right inferior fronto-occipital fasciculus (IFOF). The IFOF is a critical white matter tract responsible for higher-order cognition, serving as the primary pathway for semantic processing by connecting sensory cortices with the prefrontal lobe to enable comprehension, abstract thought, and executive control (Duffau et al., 2013). Though not statistically robust after correction, these collective findings suggest that airborne pollutants may have a subtle neurotoxic effect on the microstructural development of white matter tracts essential for executive function and attention. A reduction in neurite density within these tracts implies a structural deficit that could lower the efficiency of information transfer.

Our results could mean that no localized white-matter effects exist in this particular study population, however, they could also be taken as circumstantial evidence for a subtle effect of air pollution on white matter microstructure. Although no single association withstood correction for multiple comparisons, the consistent pattern of negative associations between pollutant exposure and neurite density across numerous tracts suggests that robust, widespread effects may not be present, but cellular-level alterations could be occurring. We hypothesize that these effects reflect changes at the cellular level, rather than broader bundle-wide alterations, a conclusion supported by the divergence of these NODDI results from our previous FBA findings (Lewandowska et al., 2025). This interpretation is biologically plausible, as exposure to air pollution has been suggested to induce neuroinflammatory processes, including microglial activation and oxidative stress (Burnor et al., 2021). It also aligns with findings from other imaging modalities that suggest air pollution can alter the cellular composition of white matter, potentially by expanding the population of glial cells (Burnor et al., 2021). The lack of statistically significant results after correction highlights the need for larger longitudinal studies to confirm these trends and elucidate the precise mechanisms through which air pollution may impact neurodevelopment.

Our study has a number of considerable strengths, which are rooted in its design approach specifically for air pollution analyses and high-quality data. We examined a population characterized by high PM exposures and large exposure ranges, which allowed for a powerful assessment of air pollution’s effects. Our exposure assessment combined comprehensive lifelong address histories with state-of-the-art air pollution models that provided high spatiotemporal resolution (see section *Supplemental Methods* in Supplementary Materials). Finally, the cohort design, which included a population of children with ADHD, along with the use of a multiple b-value diffusion MRI protocol to perform NODDI analysis, provides a comprehensive look into the complex relationship between air pollution and brain development. In addition, the analyses were prepared in the native space of each participant by registering the masks exported from the standard space to native space using T1 images. This allowed for analyses to be performed in identically defined anatomical regions across subjects. This study is not without limitations. First, as noted, the exploratory nature of the analysis with a large number of models required multiple-testing correction, which increased the threshold for statistical significance and may have led to an underestimation of the true number of affected brain regions. Second, the study did not include PM_2.5_ exposure data, a critical pollutant consistently linked to adverse health effects, which limits the comprehensiveness of our findings. While PM_10_ and PM_2.5_ are highly correlated in our study area (r = 0.9, correlation in monitoring station data) (Szwed et al., 2025), the smaller particle size of PM_2.5_ may have a stronger impact due to its greater potential for infiltrating the body’s organs, including the brain. Finally, we could not isolate the effects of specific chemical mixtures within the measured air pollutants, such as PAHs (Yang et al., 2025). While our findings link general air pollution metrics to brain changes, the specific neurotoxicants driving these associations remain unknown. A sophisticated mixture-based approach represents an important future direction but it is only recently beginning to be applied in our field of neuroscience (Bottenhorn et al., 2024).

## 5. CONCLUSIONS

The present study aimed to establish the impact of long-term PM_10_ and NO_2_ exposure across different time windows on white matter in the brains of children aged 10-13 years. Our analyses revealed several significant associations across different time windows and white matter tracts; however, no associations remained significant after correcting for multiple comparisons.

Despite the lack of robust findings, the analysis revealed numerous significant results prior to correction, suggesting that an underlying biological effect may exist, but their detection was challenged by the exploratory nature of the analysis. Consequently, these findings serve as preliminary evidence, highlighting the importance of future research focused on specific white matter regions implicated in attentional networks.

## 6. ACKNOWLEDGMENTS

We are very grateful to all of the children and their parents for their participation in the study, partner schools for helping us to reach out to children assigned to the control group, field psychologists for identifying children with ADHD and performing psychological testing on all children, and the study team for technical, logistic, administrative, and communication effort. We also thank Maja Wierzba-Łukaszyk for assistance in data acquisition.

## 7. AUTHOR CONTRIBUTION

**Paulina Lewandowska:** Conceptualization; data curation; formal analysis; visualization; writing-original draft; writing - review and editing

**Mikołaj Compa:** formal analysis

**Claude Bajada:** Conceptualization; formal analysis; writing - review and editing, supervision

**Yarema Mysak:** Data curation; software

**Aleksandra Domagalik**: Data acquisition, writing - review and editing

**Bartosz Kossowski**: Data acquisition

**Clemens Baumbach:** Data curation; software; writing - review and editing

**Katarzyna Kaczmarek-Majer:** Air pollution modelling

**Anna Degórska:** Air pollution modelling

**Krzysztof Skotak:** Air pollution modelling

**Katarzyna Sitnik-Warchulska:** Data acquisition; subject diagnosis

**Małgorzata Lipowska:** Data acquisition; subject diagnosis

**Bernadetta Izydorczyk:** Data acquisition; subject diagnosis

**James Grellier:** Conceptualization; writing – review and editing

**Iana Markevych:** Conceptualization, data curation, writing – review and editing

**Marcin Szwed:** Funding acquisition, conceptualization, data acquisition, writing - original draft; writing-review and editing, supervision

## 8. FUNDING RESOURCES

Supported by “NeuroSmog: Determining the impact of air pollution on the developing brain” (Nr. POIR.04.04.00-1763/18-00) grant to M.S. implemented as part of the TEAM-NET programme of the Foundation for Polish Science, co-financed from EU resources obtained from the European Regional Development Fund under the Smart Growth Operational Programme, by a grant from the Priority Research Area (“Anthropocene”) under the Strategic Programme Excellence Initiative at Jagiellonian University to M.S. and by a National Science Centre, Poland (grant no. K/NCN/000514) to M.S. Iana Markevych’s time on this publication was supported by the National Science Centre, Poland (grant number 2023/50/E/HS6/00102) and the “Strategic research and innovation program for the development of Medical University – Plovdiv” N◦ BG-RRP-2.004-0007-C01, Establishment of a network of research higher schools, National plan for recovery and resilience, financed by the European Union – NextGenerationEU.

## 9. DATA AVAILABILITY STATEMENT

We welcome individual data inquiries for NeuroSmog data, and engage to provide data on a case-to-case basis. Due to European Union General Data Protection Rules on sharing of sensitive data, NeuroSmog data sharing is possible on the basis of bilateral data sharing agreements ratified by the Jagiellonian University and the inquirer’s institution. Please direct inquiries to. NeuroSmog data sharing guidelines are expected to be published online in 2026 at https://neurosmog.psychologia.uj.edu.pl/dla-naukowcow/for-scientists

## REFERENCES

Andersson, J. L. R., Graham, M. S., Drobnjak, I., Zhang, H., & Campbell, J. (2018). Susceptibility-induced distortion that varies due to motion: Correction in diffusion MR without acquiring additional data. NeuroImage, 171, 277–295.

Andersson, J. L. R., Graham, M. S., Drobnjak, I., Zhang, H., Filippini, N., & Bastiani, M. (2017). Towards a comprehensive framework for movement and distortion correction of diffusion MR images: Within volume movement. NeuroImage, 152, 450–466.

Andersson, J. L. R., Graham, M. S., Zsoldos, E., & Sotiropoulos, S. N. (2016). Incorporating outlier detection and replacement into a non-parametric framework for movement and distortion correction of diffusion MR images. NeuroImage, 141, 556–572.

Andersson, J. L. R., Skare, S., & Ashburner, J. (2003). How to correct susceptibility distortions in spin-echo echo-planar images: application to diffusion tensor imaging. NeuroImage, 20(2), 870–888.

Andersson, J. L. R., & Sotiropoulos, S. N. (2016). An integrated approach to correction for off-resonance effects and subject movement in diffusion MR imaging. NeuroImage, 125, 1063– 1078.

Andersson, Jenkinson, & Smith. (2007). Non-linear registration, aka Spatial normalisation FMRIB technical report TR07JA2. FMRIB Analysis Group Of. https://www.fmrib.ox.ac.uk/datasets/techrep/tr07ja2/tr07ja2.pdf

Bastiani, M., Cottaar, M., Fitzgibbon, S. P., Suri, S., Alfaro-Almagro, F., Sotiropoulos, S. N., Jbabdi, S., & Andersson, J. L. R. (2019). Automated quality control for within and between studies diffusion MRI data using a non-parametric framework for movement and distortion correction. NeuroImage, 184, 801–812.

Bateson, T. F., & Schwartz, J. (2008). Children’s response to air pollutants. Journal of Toxicology and Environmental Health. Part A, 71(3), 238–243.

Bernanke, J., Luna, A., Chang, L., Bruno, E., Dworkin, J., & Posner, J. (2022). Structural brain measures among children with and without ADHD in the Adolescent Brain and Cognitive Development Study cohort: a cross-sectional US population-based study. The Lancet. Psychiatry, 9(3), 222–231.

Binter, A.-C., Kusters, M. S. W., van den Dries, M. A., Alonso, L., Lubczyńska, M. J., Hoek, G., White, T., Iñiguez, C., Tiemeier, H., & Guxens, M. (2022). Air pollution, white matter microstructure, and brain volumes: Periods of susceptibility from pregnancy to preadolescence. Environmental Pollution, 313, 120109.

Bottenhorn, K. L., Sukumaran, K., Cardenas-Iniguez, C., Habre, R., Schwartz, J., Chen, J.-C., & Herting, M. M. (2024). Air pollution from biomass burning disrupts early adolescent cortical microarchitecture development. Environment International, 189(108769), 108769.

Bové, H., Bongaerts, E., Slenders, E., Bijnens, E. M., Saenen, N. D., Gyselaers, W., Van Eyken, P., Plusquin, M., Roeffaers, M. B. J., Ameloot, M., & Nawrot, T. S. (2019). Ambient black carbon particles reach the fetal side of human placenta. Nature Communications, 10(1), 3866.

Burks, J. D., Bonney, P. A., Conner, A. K., Glenn, C. A., Briggs, R. G., Battiste, J. D., McCoy, T., O’Donoghue, D. L., Wu, D. H., & Sughrue, M. E. (2017). A method for safely resecting anterior butterfly gliomas: the surgical anatomy of the default mode network and the relevance of its preservation. Journal of Neurosurgery, 126(6), 1795–1811.

Burnor, E., Cserbik, D., Cotter, D. L., Palmer, C. E., Ahmadi, H., Eckel, S. P., Berhane, K., McConnell, R., Chen, J.-C., Schwartz, J., Jackson, R., & Herting, M. M. (2021). Association of Outdoor Ambient Fine Particulate Matter With Intracellular White Matter Microstructural Properties Among Children. JAMA Network Open, 4(12), e2138300.

Compa, M., Baumbach, C., Kaczmarek-Majer, K., Buczyłowska, D., Gradys, G. O., Skotak, K., Degórska, A., Bratkowski, J., Wierzba-Łukaszyk, M., Mysak, Y., Sitnik-Warchulska, K., Lipowska, M., Izydorczyk, B., Grellier, J., Asanowicz, D., Markevych, I., & Szwed, M. (2023). Air pollution and attention in Polish schoolchildren with and without ADHD. The Science of the Total Environment, 892(164759), 164759.

Connaughton, M., Whelan, R., O’Hanlon, E., & McGrath, J. (2022). White matter microstructure in children and adolescents with ADHD. NeuroImage. Clinical, 33(102957), 102957.

Cory-Slechta, D. A., Merrill, A., & Sobolewski, M. (2023). Air Pollution–Related Neurotoxicity Across the Life Span. Annual Review of Pharmacology and Toxicology, 63(1), 143–163.

Cotter, D. L., Ahmadi, H., Cardenas-Iniguez, C., Bottenhorn, K. L., Gauderman, W. J., McConnell, R., Berhane, K., Schwartz, J., Hackman, D. A., Chen, J.-C., & Herting, M. M. (2024). Exposure to multiple ambient air pollutants changes white matter microstructure during early adolescence with sex-specific differences. Communications Medicine, 4(1), 155.

Cotter, D. L., Campbell, C. E., Sukumaran, K., McConnell, R., Berhane, K., Schwartz, J., Hackman, D. A., Ahmadi, H., Chen, J.-C., & Herting, M. M. (2023). Effects of ambient fine particulates, nitrogen dioxide, and ozone on maturation of functional brain networks across early adolescence. Environment International, 177(108001), 108001.

Crooijmans, K. L. H. A., Iñiguez, C., Withworth, K. W., Estarlich, M., Lertxundi, A., Fernández-Somoano, A., Tardón, A., Ibarluzea, J., Sunyer, J., Guxens, M., & Binter, A.-C. (2024). Nitrogen dioxide exposure, attentional function, and working memory in children from 4 to 8 years: Periods of susceptibility from pregnancy to childhood. Environment International, 186(108604), 108604.

de Bont, J., Jaganathan, S., Dahlquist, M., Persson, Å., Stafoggia, M., & Ljungman, P. (2022). Ambient air pollution and cardiovascular diseases: An umbrella review of systematic reviews and meta_analyses. Journal of Internal Medicine, 291(6), 779–800.

de Hoogh, K., Wang, M., Adam, M., Badaloni, C., Beelen, R., Birk, M., Cesaroni, G., Cirach, M., Declercq, C., Dėdelė, A., Dons, E., de Nazelle, A., Eeftens, M., Eriksen, K., Eriksson, C., Fischer, P., Gražulevičienė, R., Gryparis, A., Hoffmann, B., … Hoek, G. (2013). Development of land use regression models for particle composition in twenty study areas in Europe. Environmental Science & Technology, 47(11), 5778–5786.

Duan, R.-R., Hao, K., & Yang, T. (2020). Air pollution and chronic obstructive pulmonary disease. Chronic Diseases and Translational Medicine, 6(4), 260–269.

Duffau, H., Herbet, G., & Moritz-Gasser, S. (2013). Toward a pluri-component, multimodal, and dynamic organization of the ventral semantic stream in humans: lessons from stimulation mapping in awake patients. Frontiers in Systems Neuroscience, 7, 44.

Forns, J., Dadvand, P., Foraster, M., Alvarez-Pedrerol, M., Rivas, I., López-Vicente, M., Suades-Gonzalez, E., Garcia-Esteban, R., Esnaola, M., Cirach, M., Grellier, J., Basagaña, X., Querol, X., Guxens, M., Nieuwenhuijsen, M. J., & Sunyer, J. (2016). Traffic-Related Air Pollution, Noise at School, and Behavioral Problems in Barcelona Schoolchildren: A Cross-Sectional Study. Environmental Health Perspectives, 124(4), 529–535.

Fuelscher, I., Hyde, C., Anderson, V., & Silk, T. J. (2021). White matter tract signatures of fiber density and morphology in ADHD. Cortex; a Journal Devoted to the Study of the Nervous System and Behavior, 138, 329–340.

Genc, S., Smith, R. E., Malpas, C. B., Anderson, V., Nicholson, J. M., Efron, D., Sciberras, E., Seal, M. L., & Silk, T. J. (2018). Development of white matter fibre density and morphology over childhood: A longitudinal fixel-based analysis. NeuroImage, 183, 666–676.

Global Burden of Disease Collaborative Network. Global Burden of Disease Study 2021 (GBD 2021) Air Pollution Exposure Estimates and Risk Curves 1990-2021. Seattle, United States of America: Institute for Health Metrics and Evaluation (IHME), 2024.

Guxens, M., Lubczyńska, M. J., Muetzel, R. L., Dalmau-Bueno, A., Jaddoe, V. W. V., Hoek, G., van der Lugt, A., Verhulst, F. C., White, T., Brunekreef, B., Tiemeier, H., & El Marroun, H. (2018). Air Pollution Exposure During Fetal Life, Brain Morphology, and Cognitive Function in School-Age Children. Biological Psychiatry, 84(4), 295–303.

Hartig, F., & Hartig, M. F. (2020). Package “DHARMa.” In R package. download.nust.na. http://download.nust.na/pub3/cran/web/packages/DHARMa/DHARMa.pdf

Hernandez-Fernandez, M., Reguly, I., Jbabdi, S., Giles, M., Smith, S., & Sotiropoulos, S. N. (2019). Using GPUs to accelerate computational diffusion MRI: From microstructure estimation to tractography and connectomes. NeuroImage, 188, 598–615.

Herting, M. M., Bottenhorn, K. L., & Cotter, D. L. (2024). Outdoor air pollution and brain development in childhood and adolescence. Trends in Neurosciences, 47(8), 593–607.

Herting, M. M., Younan, D., Campbell, C. E., & Chen, J.-C. (2019). Outdoor Air Pollution and Brain Structure and Function From Across Childhood to Young Adulthood: A Methodological Review of Brain MRI Studies. Frontiers in Public Health, 7, 332.

Jenkinson, M., Beckmann, C. F., Behrens, T. E. J., Woolrich, M. W., & Smith, S. M. (2012). FSL. NeuroImage, 62(2), 782–790.

Jones, D. K., Knösche, T. R., & Turner, R. (2013). White matter integrity, fiber count, and other fallacies: the do’s and don’ts of diffusion MRI. NeuroImage, 73, 239–254.

Konduracka, E., & Rostoff, P. (2022). Links between chronic exposure to outdoor air pollution and cardiovascular diseases: a review. Environmental Chemistry Letters, 20(5), 2971–2988.

Kusters, M. S. W., Binter, A.-C., Muetzel, R. L., López-Vicente, M., Petricola, S., Tiemeier, H., & Guxens, M. (2025). Outdoor residential air pollution exposure and the development of brain volumes across childhood: A longitudinal study. Environmental Pollution (Barking, Essex: 1987), 373(126078), 126078.

Kusters, M. S. W., López-Vicente, M., Muetzel, R. L., Binter, A.-C., Petricola, S., Tiemeier, H., & Guxens, M. (2024). Residential ambient air pollution exposure and the development of white matter microstructure throughout adolescence. Environmental Research, 262(Pt 2), 119828.

Lewandowska, P. (2025). Impact of prenatal and early life exposure of air pollution on NODDI metrics in Polish children: diffusion MRI study. https://osf.io/qjp2g/resources

Lewandowska, P., Bajada, C. J., Mysak, Y., Domagalik, A., Kossowski, B., Baumbach, C., Kaczmarek-Majer, K., Degórska, A., Skotak, K., Sitnik-Warchulska, K., Lipowska, M., Izydorczyk, B., Grellier, J., Markevych, I., & Szwed, M. (2025). The impact of early life exposure to air pollution on the brain: A diffusion MRI study in 10-13-year-old children with and without ADHD diagnosis. Human Brain Mapping, 46(14), e70306.

Liu, Y., Zhang, J., Rolls, E., Dai, Y., Geng, S., Deng, L., Chen, Z., Zhang, Y., Tao, M., Zhang, L., Ren, T., Feng, J., Cao, M., & Li, F. (2023). Abnormal white matter development during early childhood in autism and developmental disability. In bioRxiv. 10.1101/2023.11.15.567183

López-Vicente, M., Kusters, M., Binter, A.-C., Petricola, S., Tiemeier, H., Muetzel, R., & Guxens, M. (2025). Long-term exposure to traffic-related air pollution and noise and dynamic brain connectivity across adolescence. Environmental Health Perspectives, 133(5), 57002.

Lubczyńska, M. J., Muetzel, R. L., El Marroun, H., Basagaña, X., Strak, M., Denault, W., Jaddoe, V. W. V., Hillegers, M., Vernooij, M. W., Hoek, G., White, T., Brunekreef, B., Tiemeier, H., & Guxens, M. (2020). Exposure to Air Pollution during Pregnancy and Childhood, and White Matter Microstructure in Preadolescents. Environmental Health Perspectives, 128(2), 27005.

Lynch, K. M., Cabeen, R. P., Toga, A. W., & Clark, K. A. (2020). Magnitude and timing of major white matter tract maturation from infancy through adolescence with NODDI. NeuroImage, 212, 116672.

Mah, A., Geeraert, B., & Lebel, C. (2017). Detailing neuroanatomical development in late childhood and early adolescence using NODDI. PloS One, 12(8), e0182340.

Markevych, I., Orlov, N., Grellier, J., Kaczmarek-Majer, K., Lipowska, M., Sitnik-Warchulska, K., Mysak, Y., Baumbach, C., Wierzba-Łukaszyk, M., Soomro, M. H., Compa, M., Izydorczyk, B., Skotak, K., Degórska, A., Bratkowski, J., Kossowski, B., Domagalik, A., & Szwed, M. (2021). NeuroSmog: Determining the Impact of Air Pollution on the Developing Brain: Project Protocol. International Journal of Environmental Research and Public Health, 19(1). 10.3390/ijerph19010310

McAlonan, G. M., Cheung, V., Cheung, C., Chua, S. E., Murphy, D. G. M., Suckling, J., Tai, K.-S., Yip, L. K. C., Leung, P., & Ho, T. P. (2007). Mapping brain structure in attention deficit-hyperactivity disorder: a voxel-based MRI study of regional grey and white matter volume. Psychiatry Research, 154(2), 171–180.

Mori, S., Wakana, S., C., V. Z. P., & Nagae-Poetscher, L. M. (2005). MRI atlas of human white matter. https://books.google.com/books?hl=en&lr=&id=ltwRYlvFNLIC&oi=fnd&pg=PR5&dq=Mori+et+al.,+MRI+Atlas+of+Human+White+Matter.+Elsevier,+Amsterdam,+The+Netherlands+(2005)&ots=gfKPm98Knk&sig=27_1BeZftwLNWVXcOp5Mi7O9KKU

Morrel, J., Dong, M., Rosario, M. A., Cotter, D. L., Bottenhorn, K. L., & Herting, M. M. (2025). A systematic review of air pollution exposure and brain structure and function during development. Environmental Research, 275(121368), 121368.

Morris, R. H., Counsell, S. J., McGonnell, I. M., & Thornton, C. (2021). Early life exposure to air pollution impacts neuronal and glial cell function leading to impaired neurodevelopment. BioEssays: News and Reviews in Molecular, Cellular and Developmental Biology, e2000288.

Mortamais, M., Pujol, J., van Drooge, B. L., Macià, D., Martínez-Vilavella, G., Reynes, C., Sabatier, R., Rivas, I., Grimalt, J., Forns, J., Alvarez-Pedrerol, M., Querol, X., & Sunyer, J. (2017). Effect of exposure to polycyclic aromatic hydrocarbons on basal ganglia and attention-deficit hyperactivity disorder symptoms in primary school children. Environment International, 105, 12–19.

Nakao, T., Radua, J., Rubia, K., & Mataix-Cols, D. (2011). Gray matter volume abnormalities in ADHD: voxel-based meta-analysis exploring the effects of age and stimulant medication. The American Journal of Psychiatry, 168(11), 1154–1163.

Paciência, I., Cavaleiro Rufo, J., & Moreira, A. (2022). Environmental inequality: Air pollution and asthma in children. Pediatric Allergy and Immunology, 33(6). 10.1111/pai.13818

Parenteau, A. M., Hang, S., Swartz, J. R., Wexler, A. S., & Hostinar, C. E. (2024). Clearing the air: A systematic review of studies on air pollution and childhood brain outcomes to mobilize policy change. Developmental Cognitive Neuroscience, 69(101436), 101436.

Polemiti, E., Hese, S., Schepanski, K., Yuan, J., environMENTAL consortium, & Schumann, G. (2024). How does the macroenvironment influence brain and behaviour-a review of current status and future perspectives. Molecular Psychiatry, 1–19.

Pujol, J., Fenoll, R., Macià, D., Martínez-Vilavella, G., Alvarez-Pedrerol, M., Rivas, I., Forns, J., Deus, J., Blanco-Hinojo, L., Querol, X., & Sunyer, J. (2016). Airborne copper exposure in school environments associated with poorer motor performance and altered basal ganglia. Brain and Behavior, 6(6), e00467.

Pujol, J., Martínez-Vilavella, G., Macià, D., Fenoll, R., Alvarez-Pedrerol, M., Rivas, I., Forns, J., Blanco-Hinojo, L., Capellades, J., Querol, X., Deus, J., & Sunyer, J. (2016). Traffic pollution exposure is associated with altered brain connectivity in school children. NeuroImage, 129, 175–184.

R Core Team, R., Team, R. C., & Others. (2024). R: A language and environment for statistical computing. R Foundation for Statistical Computing, Vienna, Austria. 2012.

Simmonds, D. J., Hallquist, M. N., Asato, M., & Luna, B. (2014). Developmental stages and sex differences of white matter and behavioral development through adolescence: a longitudinal diffusion tensor imaging (DTI) study. NeuroImage, 92, 356–368.

Skandalakis, G. P., Komaitis, S., Kalyvas, A., Lani, E., Kontrafouri, C., Drosos, E., Liakos, F., Piagkou, M., Placantonakis, D. G., Golfinos, J. G., Fountas, K. N., Kapsalaki, E. Z., Hadjipanayis, C. G., Stranjalis, G., & Koutsarnakis, C. (2020). Dissecting the default mode network: direct structural evidence on the morphology and axonal connectivity of the fifth component of the cingulum bundle. Journal of Neurosurgery, 134(4), 1–12.

Stave, E. A., De Bellis, M. D., Hooper, S. R., Woolley, D. P., Chang, S. K., & Chen, S. D. (2017). Dimensions of attention associated with the microstructure of corona radiata white matter. Journal of Child Neurology, 32(5), 458–466.

Sunyer, J., Esnaola, M., Alvarez-Pedrerol, M., Forns, J., Rivas, I., López-Vicente, M., Suades-González, E., Foraster, M., Garcia-Esteban, R., Basagaña, X., Viana, M., Cirach, M., Moreno, T., Alastuey, A., Sebastian-Galles, N., Nieuwenhuijsen, M., & Querol, X. (2015). Association between traffic-related air pollution in schools and cognitive development in primary school children: a prospective cohort study. PLoS Medicine, 12(3), e1001792.

Szwed, M., de Jesus, A. V., Kossowski, B., Ahmadi, H., Rutkowska, E., Mysak, Y., Baumbach, C., Kaczmarek-Majer, K., Degórska, A., Skotak, K., Sitnik-Warchulska, K., Lipowska, M., Grellier, J., Markevych, I., & Herting, M. M. (2025). Air pollution and cortical myelin T1w/T2w ratio estimates in school-age children from the ABCD and NeuroSmog studies. Developmental Cognitive Neuroscience, 73(101538), 101538.

Takahashi, M., Iwamoto, K., Fukatsu, H., Naganawa, S., Iidaka, T., & Ozaki, N. (2010). White matter microstructure of the cingulum and cerebellar peduncle is related to sustained attention and working memory: a diffusion tensor imaging study. Neuroscience Letters, 477(2), 72–76.

Thygesen, M., Holst, G. J., Hansen, B., Geels, C., Kalkbrenner, A., Schendel, D., Brandt, J., Pedersen, C. B., & Dalsgaard, S. (2020). Exposure to air pollution in early childhood and the association with Attention-Deficit Hyperactivity Disorder. Environmental Research, 183(108930), 108930.

Torchiano, B., & Wheeler, M. (2025). Package “lmPerm.” In R package version. mirror.las.iastate.edu. https://mirror.las.iastate.edu/CRAN/web/packages/lmPerm/lmPerm.pdf

Vaher, K., Galdi, P., Blesa Cabez, M., Sullivan, G., Stoye, D. Q., Quigley, A. J., Thrippleton, M. J., Bogaert, D., Bastin, M. E., Cox, S. R., & Boardman, J. P. (2022). General factors of white matter microstructure from DTI and NODDI in the developing brain. NeuroImage, 254(119169), 119169.

Yang, H., Cohen, J. W., Pagliaccio, D., Ramphal, B., Rauh, V., Perera, F., Peterson, B. S., Andrews, H., Rundle, A. G., Herbstman, J., & Margolis, A. E. (2025). Prenatal exposure to polycyclic aromatic hydrocarbons, reduced hippocampal subfield volumes, and word reading. Developmental Cognitive Neuroscience, 72(101508), 101508.

Zundel, C. G., Ely, S., Brokamp, C., Strawn, J. R., Jovanovic, T., Ryan, P., & Marusak, H. A. (2024). Particulate matter exposure and default mode network equilibrium during early adolescence. Brain Connectivity, 14(6), 307–318.

